# Identifying Ecological Restoration Priority Areas through a Function–Structure–Dynamic Framework: Integrating Ecological Security Patterns and Future Land-Use Simulation in the Anhui Section of the Yangtze River Basin

**DOI:** 10.64898/2026.06.13.732028

**Authors:** Yuhan Xia, Shanshan Liang, Ping Mi, Ziyu Chen, Yiwen Fan, Wenjing Zhao, Ruonan Li

## Abstract

Rapid urbanization has intensified ecological degradation and threatened ecosystem sustainability in the Yangtze River Basin. This study proposes a Function–Structure–Dynamic (FSD) framework integrating ecological protection importance assessment, ecological security pattern (ESP) construction, and future land-use simulation to identify ecological restoration priorities in the Anhui section of the Yangtze River Basin. Ecologically important areas were identified by coupling ecosystem services and ecological vulnerability. MSPA, landscape connectivity analysis, resistance surface modeling, and circuit theory were applied to construct the ESP, while the PLUS model simulated land-use change and ecological risks in 2032. Results identified 37 ecological sources covering 6,901.39 km² and 84 ecological corridors with a total length of 916.80 km. Ecological pinch points, barrier areas, and 17 ecological warning areas were further delineated. Based on these findings, four ecological restoration zones were proposed with differentiated management strategies. The FSD framework effectively integrates ecological functions, spatial structure, and future dynamics, providing scientific support for ecological restoration and sustainable territorial spatial planning in rapidly urbanizing regions.

## 1. Introduction

With rapid socioeconomic development and accelerating urbanization, high-intensity land development has undermined ecosystem stability and posed significant threats to regional ecological security and sustainable development (1, 2). In response, national policies emphasize integrated natural resource planning, the enhancement of ecosystem diversity, stability, and sustainability, and the promotion of ecological protection and restoration (3). Ecological restoration has increasingly shifted toward a global, systematic, and science-based approach, creating an urgent need to identify priority restoration areas through comprehensive spatial analysis. As an effective framework for identifying priority conservation areas, guiding socioeconomic development, and balancing ecological protection, the ecological security pattern has become a core national strategy for achieving high-level ecological security and regional sustainable development (4–6).

At present, the ecological security pattern (ESP) has emerged as an important research framework, commonly constructed using the “source–resistance surface–corridor–node” paradigm (7–9). The identification of ecological sources is primarily based on patch morphological characteristics and ecological importance (10, 11). Patch morphological characteristics are commonly assessed using morphological spatial pattern analysis (MSPA) (12, 13), landscape connectivity analysis (14), and granularity backstepping methods (15). Patch ecological importance is generally evaluated based on ecosystem service function assessment (8, 16), ecosystem service supply–demand relationships (17, 18), and ecosystem service valuation methods (19, 20).

Ecological resistance surfaces are constructed by integrating multiple evaluation factors into different index systems. The analytic hierarchy process (AHP) and expert consultation methods are commonly used to assign resistance weights; however, these approaches involve a high degree of subjectivity (21, 22). At present, no unified standard exists for resistance factor selection, and regional heterogeneity must be considered to ensure the accurate application of resistance surfaces (23). Ecological corridors are typically identified using the minimum cumulative resistance (MCR) model (24, 25) and circuit theory (26). Compared with the MCR model, circuit theory can identify multiple potential corridors with spatial width, providing greater practical applicability (26, 27).Domestic ecological security assessments have mainly focused on ecological carrying capacity, ecological risk, and ecological security barriers, with an emphasis on quantifying landscape connectivity changes in key areas. However, the identification of ecological corridors, pinch points, and barrier areas remains largely conceptual, with limited spatial implementation. Therefore, stronger integration between the protection of ecological priority areas and spatial planning is required to promote the transition of ecological security patterns from theoretical frameworks to practical applications.

The Anhui section of the Yangtze River Basin includes Anqing, Chizhou, Tongling, Wuhu, and Ma’anshan and is located in a key region of the middle and lower reaches of the Yangtze River. The region has a complex and diverse geographical environment and performs critical ecological functions, including water conservation and soil and habitat maintenance, serving as an important ecological security barrier (28). However, driven by urban expansion and industrial development, ecological problems such as wetland shrinkage, excessive shoreline development, water pollution, and vegetation fragmentation have become increasingly prominent (29). Although various measures have been implemented to address these issues and have achieved notable progress, limitations remain due to insufficient integration of economic development and ecological restoration, which has continued to affect ecosystem stability and service functions (30). Therefore, there is an urgent need to scientifically identify priority areas for ecological restoration and implement differentiated restoration strategies within the framework of territorial spatial planning, making this a critical issue for ecological governance and sustainable development (4, 31).

In view of the above, this study aims to: (1) Incorporate ecological corridor width into the construction of the ecological security pattern (ESP) and determine the ESP of the study area. (2) Address the region’s key ecological problems by developing an Importance of Ecological Protection (IEP) evaluation system based on the ESP, in order to identify priority ecological protection areas. (3) Analyze the coupling relationship between construction land expansion and eco-environmental effects in 2032, identify ecological warning areas with potential risks, and delineate key ecological areas for 2032. (4) Predict ecological planning zones for the study area in 2032 and propose differentiated management strategies to provide a scientific basis for future territorial spatial planning.

## 2. Materials and methods

### 2.1 Study area

The Anhui section of the Yangtze River Basin is located in eastern China, within Anhui Province. It lies along the lower reaches of the Yangtze River, extending from 29°56′–32°06′ N and 115°75′–118°88′ E. The study area extends from Bowang District of Ma’anshan City in the east to Taihu County of Anqing City in the west, and from Ma’anshan City and County in the north to Chizhou City in the south. Covering approximately 34,919 km², the region includes five prefecture-level cities and 31 counties (urban districts), namely Anqing, Chizhou, Tongling, Wuhu, and Ma’anshan (Fig. 1). The region features a diverse physical geography, characterized by the coexistence of mountains, hills, plains, and dense water networks, and includes prominent natural landscapes such as the Yangtze River and the Dabie Mountains. It has a distinct four-season climate with abundant precipitation, diverse and fertile soils, sufficient water resources, high forest coverage, rich biodiversity, and abundant natural resources.

**Fig. 1.**
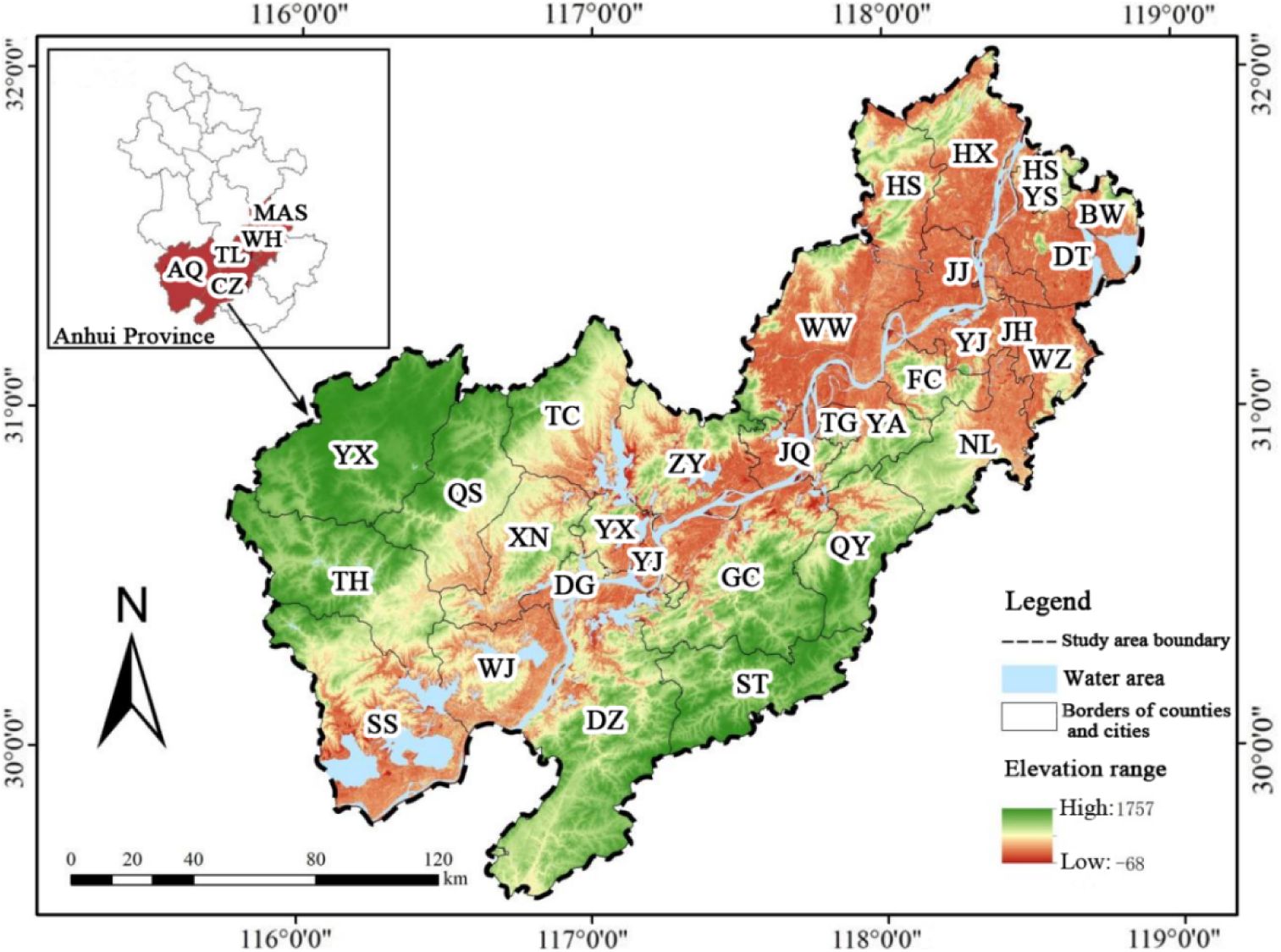
Overview of the study area.

As a key region for economic development, agricultural production, and ecological conservation in Anhui Province, the area is under increasing pressure from rapid population growth and economic expansion. By 2023, the resident population had reached 130.56 million, accompanied by accelerated development and intensified population concentration, leading to wetland degradation, excessive shoreline development, worsening water pollution, and increased vegetation fragmentation, which undermine ecosystem stability and ecosystem service functions. Consequently, the scientific identification of priority ecological restoration areas, the prediction of future ESP, and the development of differentiated restoration strategies have become critical tasks for ecological governance and sustainable development.

### 2.2 Data sources

The data used in this study were obtained from the following sources.

(1) Land use data for 2017 and 2022, with a spatial resolution of 300 m, were obtained from the European Space Agency (ESA) Climate Change Initiative (CCI) dataset (http://maps.elie.ucl.ac.be/CCI/viewer/). The land use data were reclassified into six landscape types—forest land, grassland, water area, cropland, construction land, and unused land—using ArcGIS Pro.

(2) Digital elevation model (DEM) data with a spatial resolution of 30m were obtained from the Geospatial Data Cloud (https://www.gscloud.cn/search).

(3) Soil and vegetation data were obtained from the FAO World Soil Database (1km resolution; https://www.fao.org/). The normalized difference vegetation index (NDVI) data (1km resolution) were obtained from the Earth Resources Data Cloud Platform (http://www.gis5g.com/). Net primary productivity (NPP) data (500m resolution) were derived from the MODIS dataset (https://pdaac.vsgs.gov/product_search/). Root-restricting layer depth data were obtained from the Global Change Research Data Publishing Platform (http://globalchange.bnu.edu.cn/research/cdtb.jsp).

(4) Traffic network data were obtained from the Geospatial Data Cloud (https://www.gscloud.cn/search) and processed using Euclidean distance analysis.

(5) Socio-economic data were obtained from the Anhui Provincial Statistical Yearbook and the statistical yearbooks of the respective cities.

All datasets were processed using ArcGIS 10.8 and projected to the coordinate system (WGS 1984 UTM Zone 50N).

### 2.3 Methods

This study adopts the framework model of “Importance of Ecological Protection (IEP) Evaluation—Ecological Security Pattern (ESP) Construction—Key Area Identification—Prediction of Differentiated Restoration Zoning”. It aims to predict ecologically critical areas in 2032 and propose spatial planning and restoration strategies for the region. The research framework is shown in Fig 2.

**Fig. 2.**
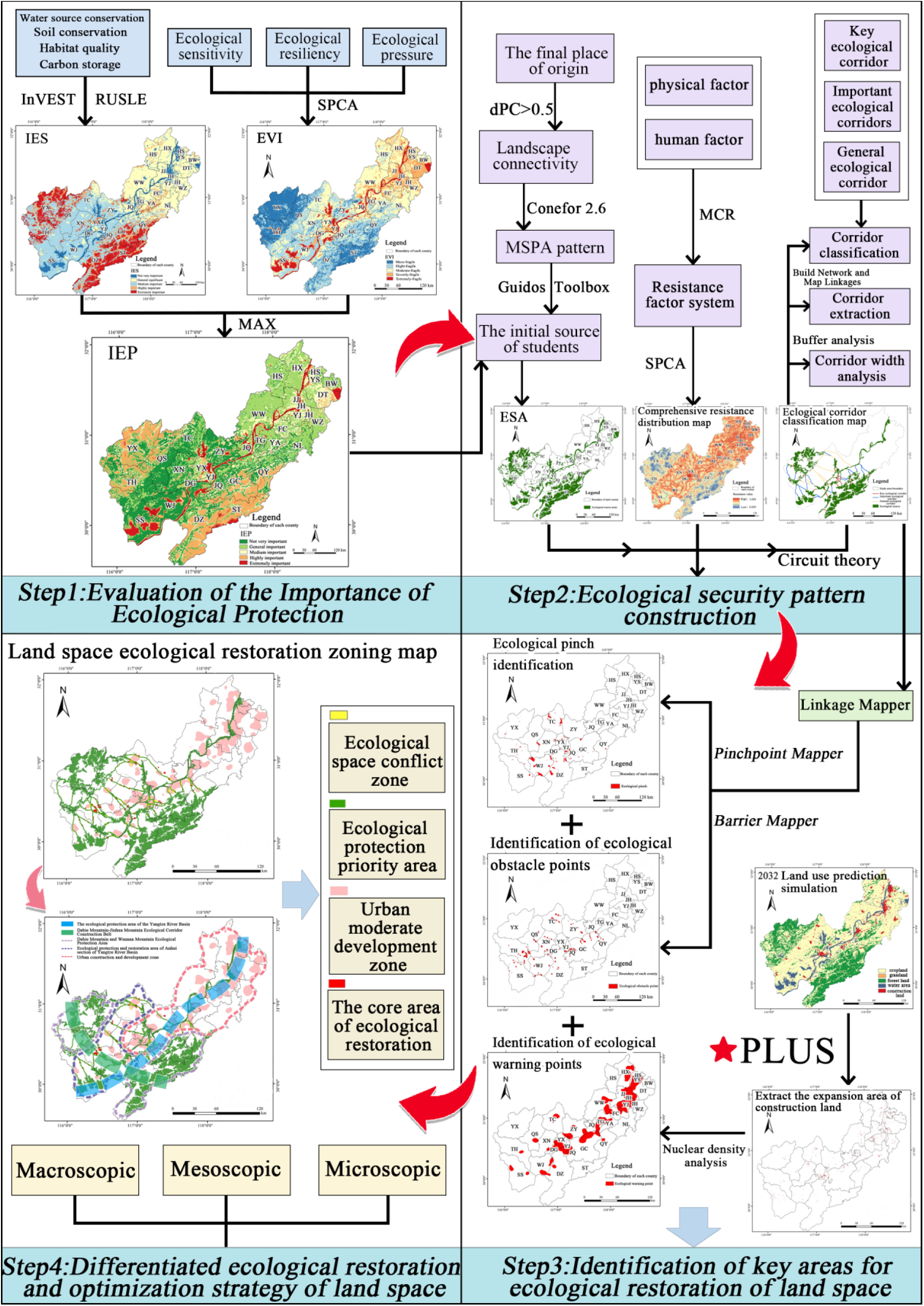
The research framework.

#### 2.3.1 Evaluation of the Importance of Ecological Protection

The importance of ecosystem services and the results of ecological vulnerability assessment were overlaid, and the maximum value method was applied to evaluate the Importance of Ecological Protection (IEP), thereby identifying priority ecological protection areas in the Anhui section of the Yangtze River Basin. The calculation formula is as follows:

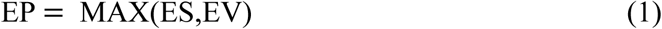

Where EP represents the Importance of Ecological Protection; ES denotes the Importance of Ecosystem Services; EV refers to the Ecological Vulnerability Index.

##### (1) Assessment of the importance of ecosystem services

Ecosystem service functions refer to the contributions of ecosystems to human production and well-being (16). Based on the key ecological problems of the study area, this study focuses on ecosystem services that support human survival and quality of life. Using spatial principal component analysis (SPCA), four ecosystem services that best represent the study area were selected that water conservation, soil conservation, habitat quality, and carbon storage. These services were combined using an equal-weight overlay to assess their spatial distribution, yielding an integrated evaluation of ecosystem service importance (Tab. 1).

**Tab. 1.**
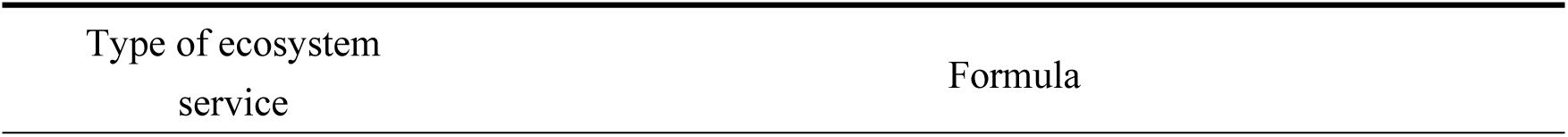
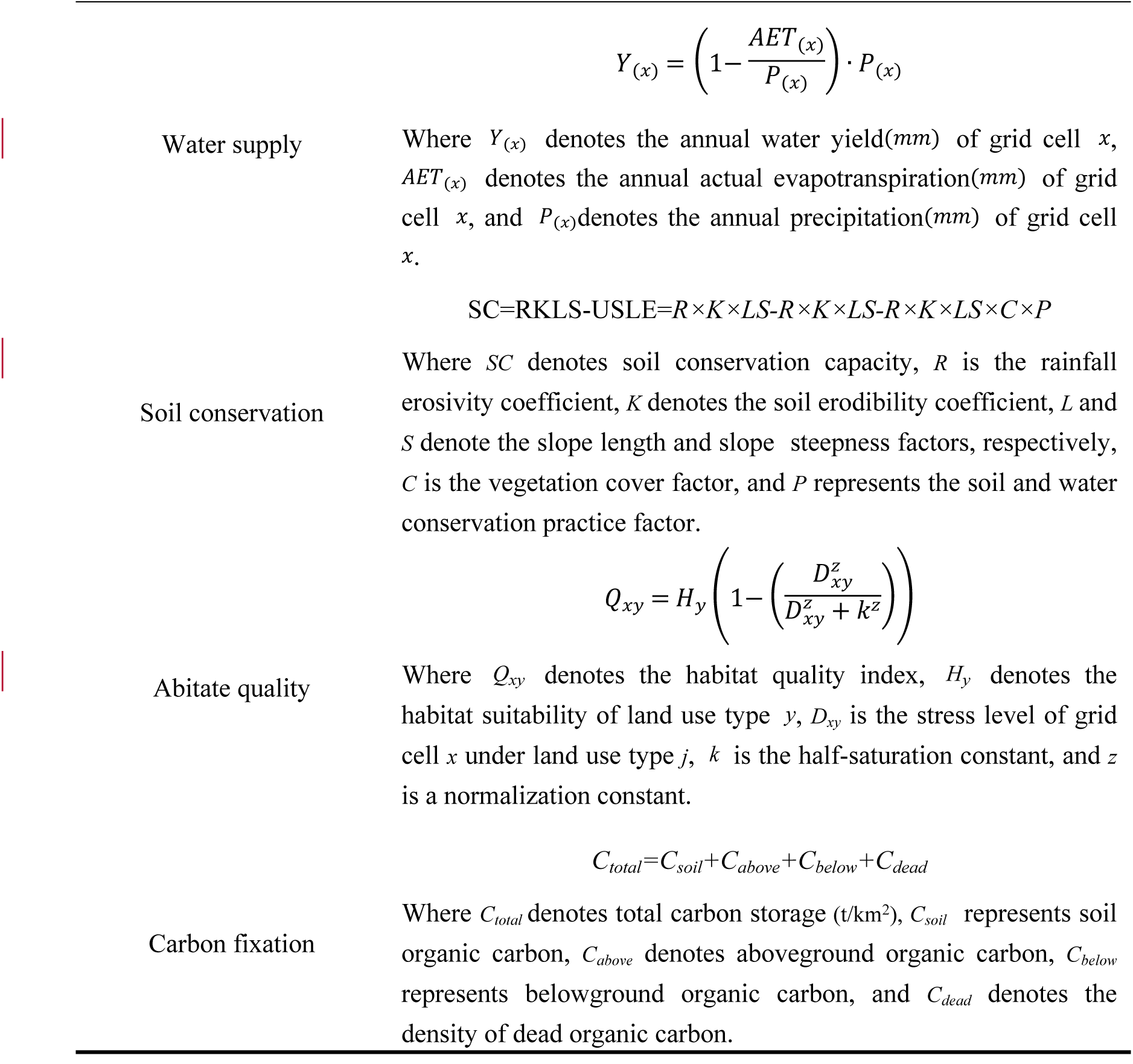
Assessment model of ecosystem service functions.

##### (2) Ecological vulnerability assessment system

Based on the Sensitivity–Recovery–Pressure (SRP) model, an ecological vulnerability assessment system was established to conduct a comprehensive evaluation from three dimensions: ecological sensitivity, ecological resilience, and ecological pressure (32). Spatial principal component analysis (SPCA) was applied to determine the weights of the indicators. Both positive and negative indicators were standardized, and all indicator values were normalized to a range of 0–1 (33). The calculation formula for the ecological vulnerability index (EVI) under the SRP model is as follows:

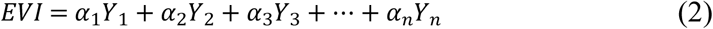

Where *EVI* denotes the ecological vulnerability index, *α*_*n*_ represents the contribution rate of the *n*th principal component, *Y*_*n*_ denotes the *n*th principal component score, and (*n*= 5).

##### (3) Landscape connectivity evaluation

Landscape connectivity is a key indicator for assessing the integrity of ecological networks and the feasibility of species migration, and it is widely used to evaluate landscape structure and function (34). The patch importance index (dPC) serves as a direct measure of landscape connectivity and is commonly applied to identify the relative importance of habitat patches. Based on previous studies, patches with a dPC value greater than 0.5 in the preliminarily screened ecological sources were selected as final ecological sources. This paper selects three indexes : IIC, PC and dPC. The relevant formulas are as follows :

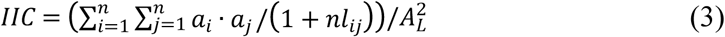

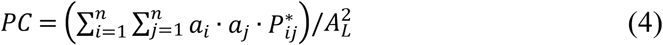

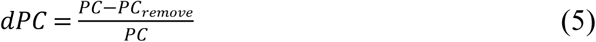

Where *n* represents the total number of patches, *a*_*i*_ and *a*_*j*_ denote the area of patch *i* and patch *j*, respectively, *A*_*L*_ is the total landscape area; *nl*_*ij*_ refers to the number of patches along the shortest path between *i* and *j*, 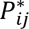 denotes the maximum probability of species dispersal between patches *i* and *j*; *PC* is the probability of connectivity index; and *PC*_*remove*;_ represents the overall probability of connectivity of the remaining patches. A larger *dPC* value indicates stronger landscape connectivity and higher ecological importance of the corresponding patch.

#### 2.3.2 Construction of comprehensive resistance surface

Based on existing studies and the local conditions of the study area, six core resistance factors, including elevation, slope, land use type, fractional vegetation cover (FVC), distance to water area, and distance to roads, were selected to construct the comprehensive resistance surface (10, 35). Using the natural breaks method, each resistance factor was classified into five levels (Tab. 2), with higher resistance values corresponding to higher resistance levels. The weights of the resistance factors were determined using spatial principal component analysis (SPCA), based on which both individual resistance surfaces and an integrated resistance surface were constructed.

**Tab. 2.** Resistance factor and grade division.

| Index factor | Resistance assignment |  |  |  |  |
| --- | --- | --- | --- | --- | --- |
|  | 1 | 2 | 3 | 4 | 5 |
| Elevation(m) | <111 | 111-302 | 302-542 | 542-866 | >866 |
| Slope(°) | <4.80 | 4.80-11.77 | 11.77-20.47 | 20.47-30.61 | >30.61 |
| Land use type | Forest land,<br>wetland and<br>water area | Grassland | Cropland | Unused land | Construction<br>land |
| FVC | >0.86 | 0.76-0.86 | 0.60-0.76 | 0.34-0.60 | <0.34 |
| Distance from<br>water system(m) | <200 | 200-500 | 500-1000 | 1000-2000 | >2000 |
| Distance from<br>road(m) | >5000 | 3000-5000 | 1500-3000 | 500-1500 | <500 |

#### 2.3.3 Ecological corridor and suitable width, key node identification

Circuit theory was used to simulate cumulative current flow between ecological sources, thereby identifying potential ecological corridors and key nodes with explicit width information (36). Based on the optimized resistance threshold, the least-cost path (LCP) method was further applied to determine the optimal spatial configuration of ecological corridors (37).Instead of applying a fixed empirical threshold, corridor width was optimized by evaluating the sensitivity of cumulative current density and the proportion of non-ecological land across multiple resistance thresholds. This data-driven approach improves the objectivity and reproducibility of corridor width selection and enhances its applicability to spatial planning (38).

Ecological pinch points and ecological obstacle points are critical nodes within ecological corridors. Ecological pinch points represent key locations that connect multiple ecological regions or species habitats, and areas with high current density indicate priority locations for maintaining connectivity (39). In contrast, ecological obstacle points impede species movement and gene flow, and their removal or mitigation can significantly enhance landscape connectivity and reduce ecological resistance (1, 40). Accordingly, both pinch points and obstacle points were identified as priority areas for ecological restoration.

#### 2.3.4 PLUS model predicts ecological planning in 2032

Ecological warning points are used to predict areas at risk of ecological degradation induced by future spatial development (41). The PLUS model, integrating the land expansion analysis strategy (LEAS) and a cellular automata model based on multi-type random patch seeds (CARS), was employed to quantify the contribution of driving factors and to simulate spatiotemporal land use change dynamics (42). Using land use data from 2017 and 2022, ecological sources were set as restricted development areas. Based on previous studies (43), ten driving factors were selected, including elevation, slope, mean annual precipitation, mean annual temperature, population density, per capita GDP, distance to railways, distance to highways, distance to water area, and distance to residential areas, to simulate construction land expansion from 2022 to 2032. Kernel density analysis was applied to identify hotspots of construction land expansion, which were overlaid with the ecological network to determine the spatial distribution of ecological warning points. Consistent with existing research, ecological warning points were closely associated with construction land expansion (44). Based on these results, key areas for ecological restoration in 2032 were identified to support future ecological planning. Model validation showed that the simulation of 2022 land use based on 2017 data achieved a Kappa coefficient of 0.96, indicating high accuracy.

## 3. Results

### 3.1 Distribution of ecological sources

#### 3.1.1 Evaluation of the importance of ecological protection (IEP)

The spatial pattern of ecological protection importance in the study area is characterized by river-oriented continuity, mountain dominance, wetland mosaics, and urban fragmentation (Fig. 3). The ecological protection barrier exhibits a spatial configuration of “two mountains and one river.” Areas of extremely high ecological protection importance are primarily composed of wetland and water area and are concentrated along the urban belt of the Yangtze River, with additional clusters in Susong and Wangjiang. Areas of high ecological protection importance are mainly forest land, distributed across the Dabie Mountains and Jiuhua Mountains, including Yuexi, Taihu, Qianshan, Shitai, Dongzhi, Guichi, and Qingyang. In contrast, areas with low ecological protection importance are mainly located in the central and western parts of the study area, dominated by construction land and strongly influenced by human activities.

**Fig. 3.**
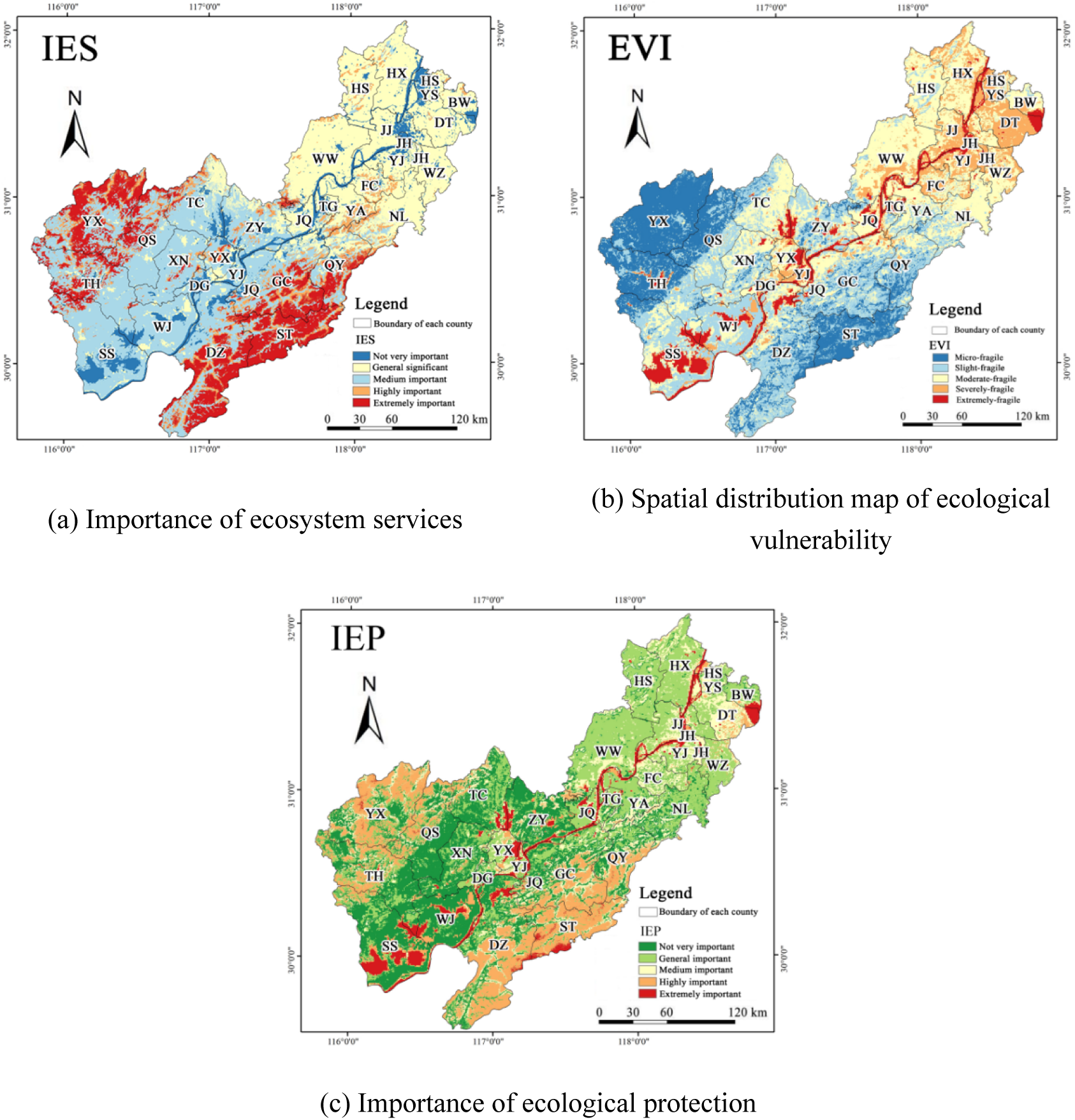
The evaluation results of the importance of ecological protection.

#### 3.1.2 Ecological source identification

Considering the scale of the study area, patches classified as extremely important and highly important in ecological protection importance were extracted as initial ecological sources (Fig. 4a). Using Conefor 2.6 software, the connection distance threshold was set to 2000 m and the connectivity probability was set to 0.5. Based on the patch importance index (dPC), 37 patches with dPC values greater than 0.5 were selected as final ecological sources (Fig. 4b). These sources cover an area of 6,901.39 km², accounting for 19.76% of the total study area, and are mainly composed of forest land, wetland, and water area. Spatially, they are concentrated in Yuexi, Wangjiang, Susong, and Taihu in Anqing, as well as Dongzhi, Guichi, Shitai, and Qingyang in Chizhou. Overall, the ecological sources exhibit a spatial pattern of “two mountains and one river,” corresponding to the Dabie Mountains, Jiuhua Mountains, and the Yangtze River corridor. Together, mountain forests, wetland and water systems, and riparian green spaces form the core ecological security pattern of the Anhui section of the Yangtze River Basin.

**Fig. 4.**
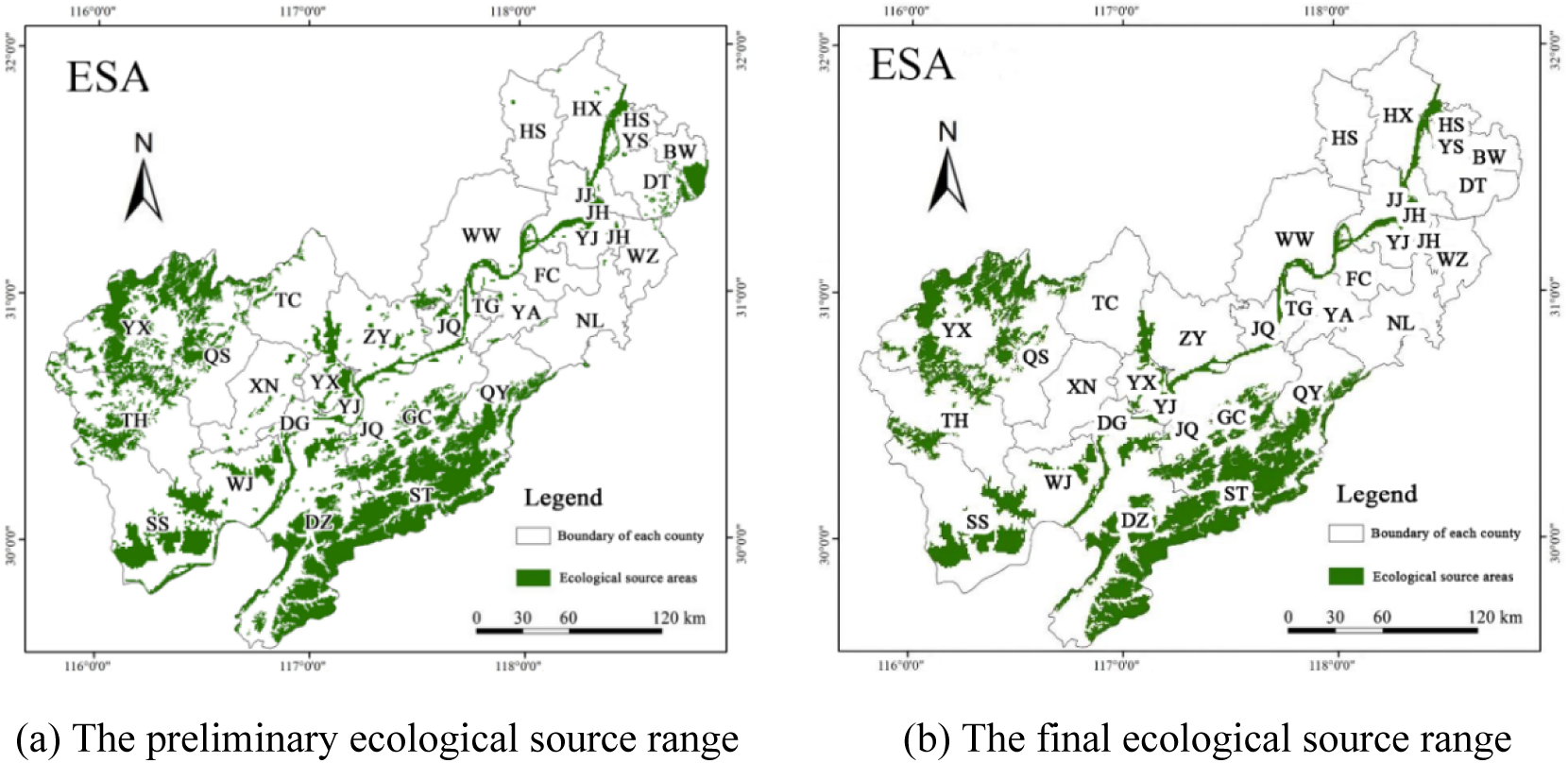
The distribution of ecological sources in the study area.

### 3.2 Construction of ecological resistance surface

Based on the relative weights of resistance factors, a single resistance surface was generated using the grid calculator, and a comprehensive resistance surface was subsequently obtained through spatial principal component analysis (SPCA). The comprehensive resistance surface exhibits a spatial pattern characterized by high resistance in urban centers and a gradual decrease toward surrounding areas (Fig. 5). High-resistance zones are mainly concentrated in densely populated and economically developed urban areas, particularly in Wuhu and Ma’anshan, where construction land dominates and expansion occurs along major transportation corridors. In contrast, low-resistance zones are primarily distributed in mountainous forest land, wetland, and water area in Yuexi, Qianshan, and Shitai. Resistance values in mountainous areas are slightly lower than those in river and lake basins, reflecting limited human disturbance, high habitat quality, and favorable conditions for species habitation and reproduction.

**Fig. 5.**
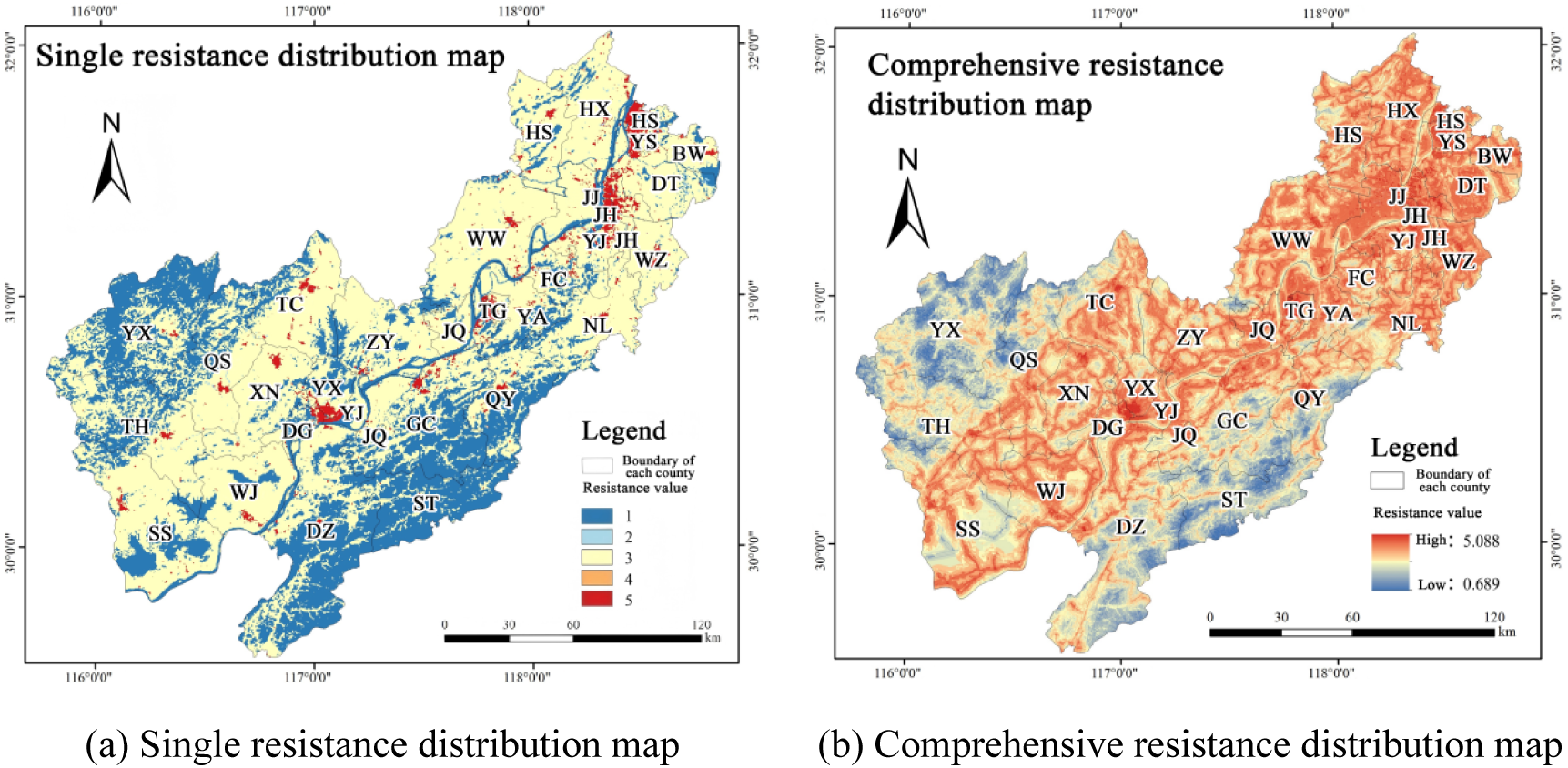
The distribution map of single and comprehensive resistance surface in the study area.

### 3.3 Identification of important ecological regions

#### 3.3.1 Identification of ecological corridor

A total of 84 ecological corridors were identified, with a combined length of 916.80 km and an average length of 10.91 km. The spatial distribution of corridors exhibits pronounced heterogeneity, characterized by higher density in the west than in the east and stronger connectivity in the south than in the north. Among them, 57 key ecological corridors, with a total length of 121.41 km, are mainly concentrated in the Dabie Mountain and Jiuhua Mountain, where ecological resources are abundant. Nineteen important ecological corridors, totaling 402.63 km, primarily link the western and southeastern mountainous areas with ecological sources along the river basin, thereby enhancing landscape connectivity between the eastern and western parts of the study area. In addition, eight general ecological corridors, with a total length of 392.76 km, are mainly distributed in the central and western regions. These corridors largely traverse areas of intensive human activity and serve as connecting pathways between ecological sources in the Dabie Mountain and Tianzhu Mountain. Although relatively long due to the limited number of large ecological sources in the central river basin, these corridors play a critical role in maintaining regional ecological connectivity and facilitating species movement (Fig. 1).

**Fig. 2.**
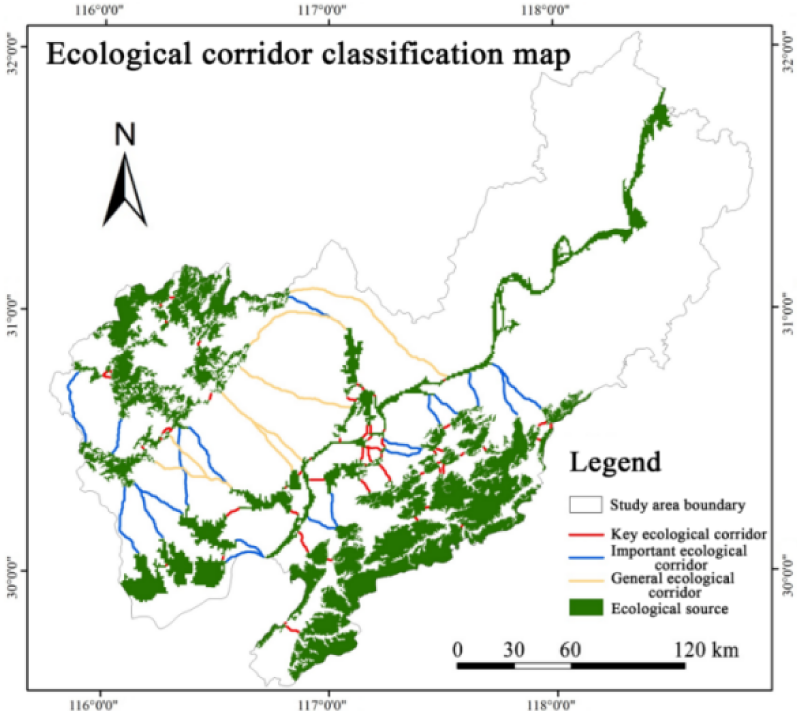
Ecological corridor classification map from 2002 to 2022.

Cropland is the main coverage type of ecological corridor, followed by forest land. The proportion of cropland and construction land decreases with the decrease of width. The proportion of woodland, wetland and water and grassland increased. It is known that the expansion of construction land will hinder energy flow. Considering the factors of ecological function, land use, economic cost and human activities, it is suggested that 30-100 m wide ecological corridor is the best choice (Fig. 7).

**Fig. 7.**
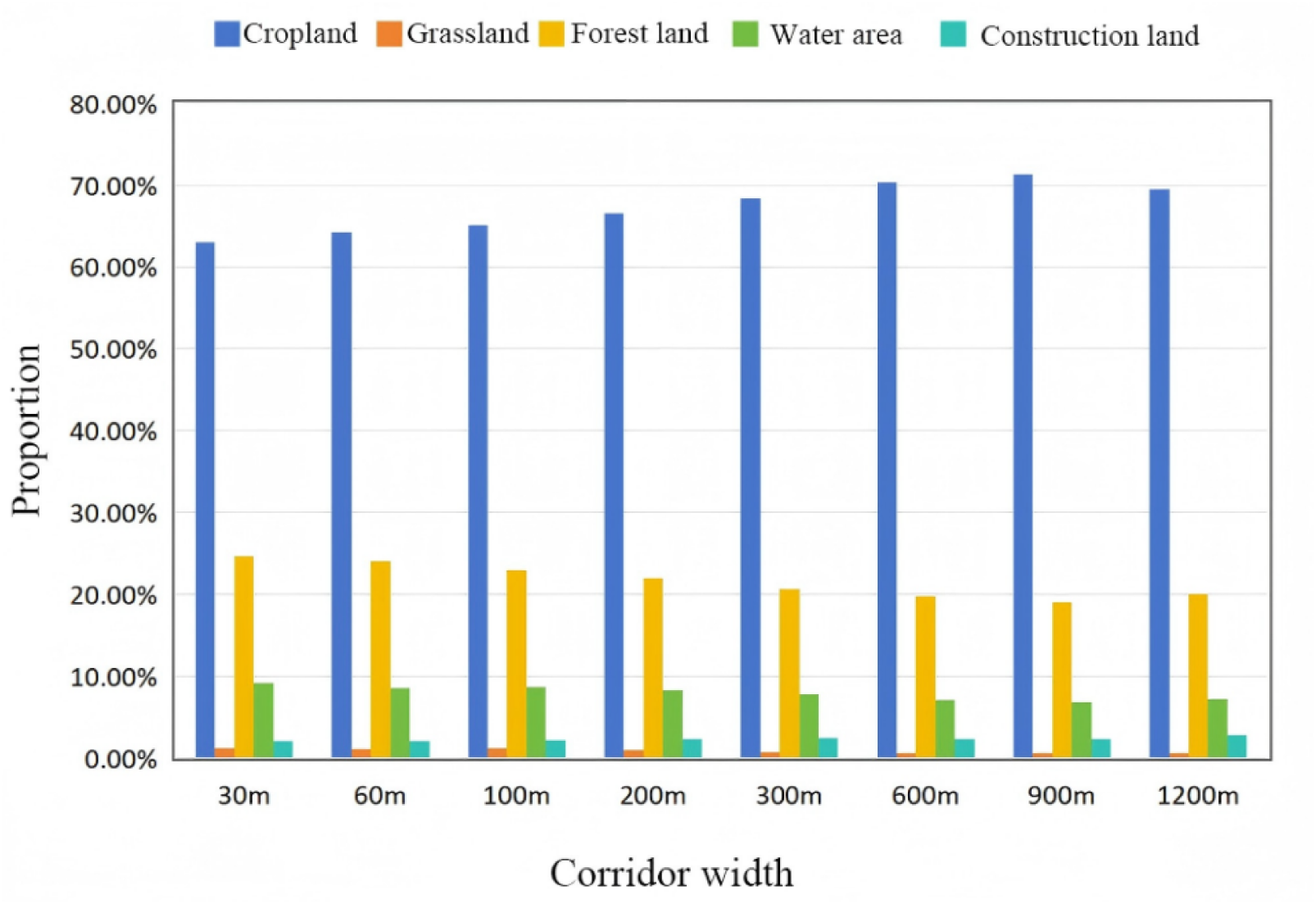
Composition of land use types of ecological corridors with different widths.

#### 3.3.2 Ecological pinch identification

Ecological pinch points in the study area cover a total area of 419.64 km². Spatially, they exhibit a heterogeneous distribution pattern characterized by higher density in the south than in the north and in the west than in the east. Ecological pinch points are mainly concentrated in Taihu, Susong, Wangjiang, and Huaining counties in Anqing, as well as Dongzhi, Guichi, and Qingyang in Chizhou. These areas are critical for maintaining overall ecological stability and therefore require targeted protection and management. Land use analysis indicates that ecological pinch points are predominantly located on cropland (66.16%), followed by forest land (21.11%), suggesting that human activities, particularly urbanization and agricultural intensification, have a substantial influence on ecological connectivity. This pattern reflects the coexistence of urban and agricultural land uses in the study area and provides an important basis for regional ecological protection and subsequent spatial planning (Fig. 8).

**Fig. 8.**
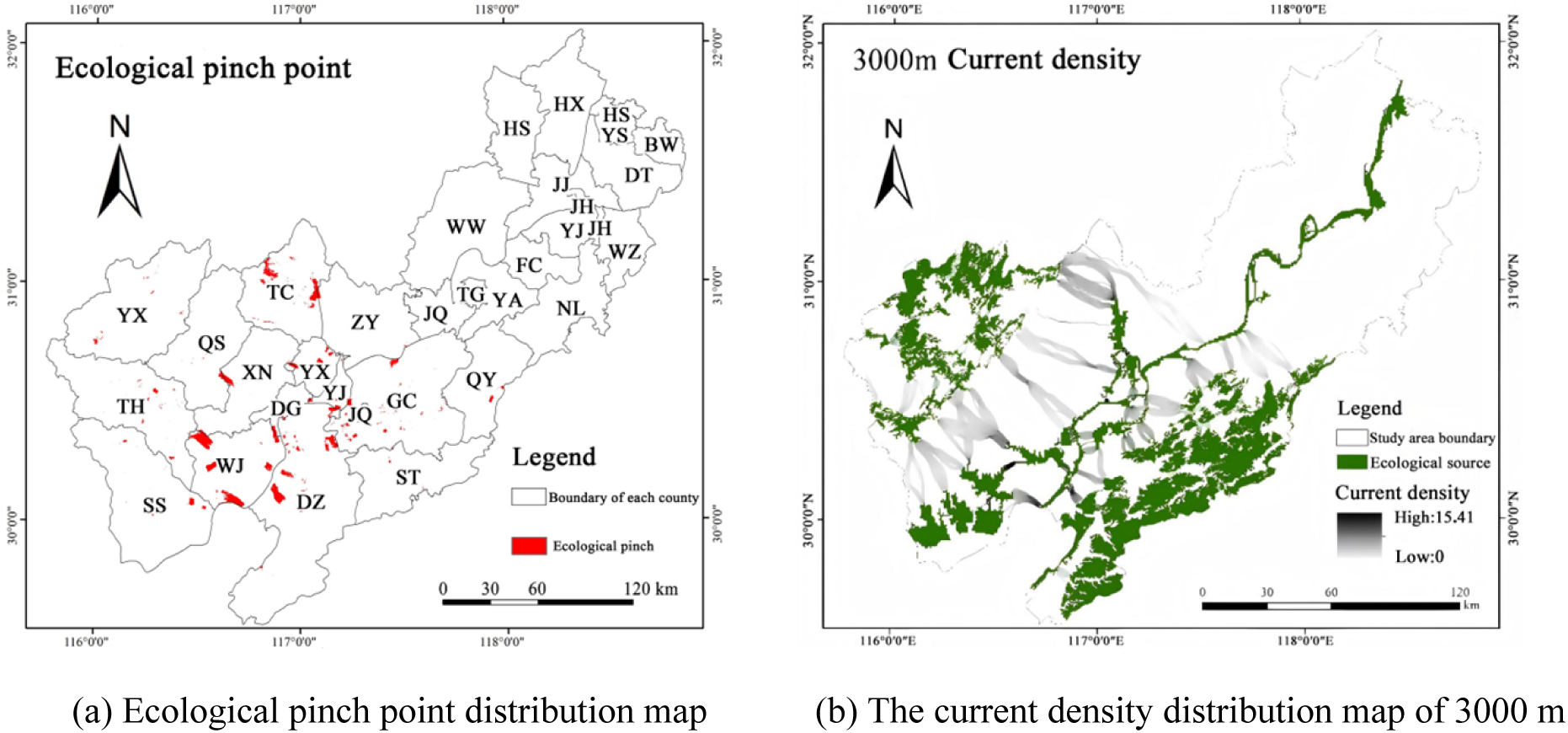
The distribution map of ecological obstacle points and 3000 m current density.

#### 3.3.3 Identification of ecological obstacle points

Ecological obstacle points cover a total area of 515.81 km², of which 418.16 km² (81.07%) are concentrated in patches larger than 30 km². Spatially, these obstacle points are mainly distributed in Anqing and Chizhou and exhibit a pattern of higher density in the eastern and western regions and lower density in the central area. In terms of land use composition, cropland dominates ecological obstacle points, accounting for 83.83% of the total area. Furthermore, ecological obstacle points are relatively concentrated in mountainous and hilly areas, where ecological corridor connectivity is severely constrained. This pattern indicates accelerated urbanization in these regions, as population growth, infrastructure development, and industrial expansion have driven the conversion of land toward construction land, intensifying the spatial squeeze effects of urbanization and industrialization (Fig. 9).

**Fig. 9.**
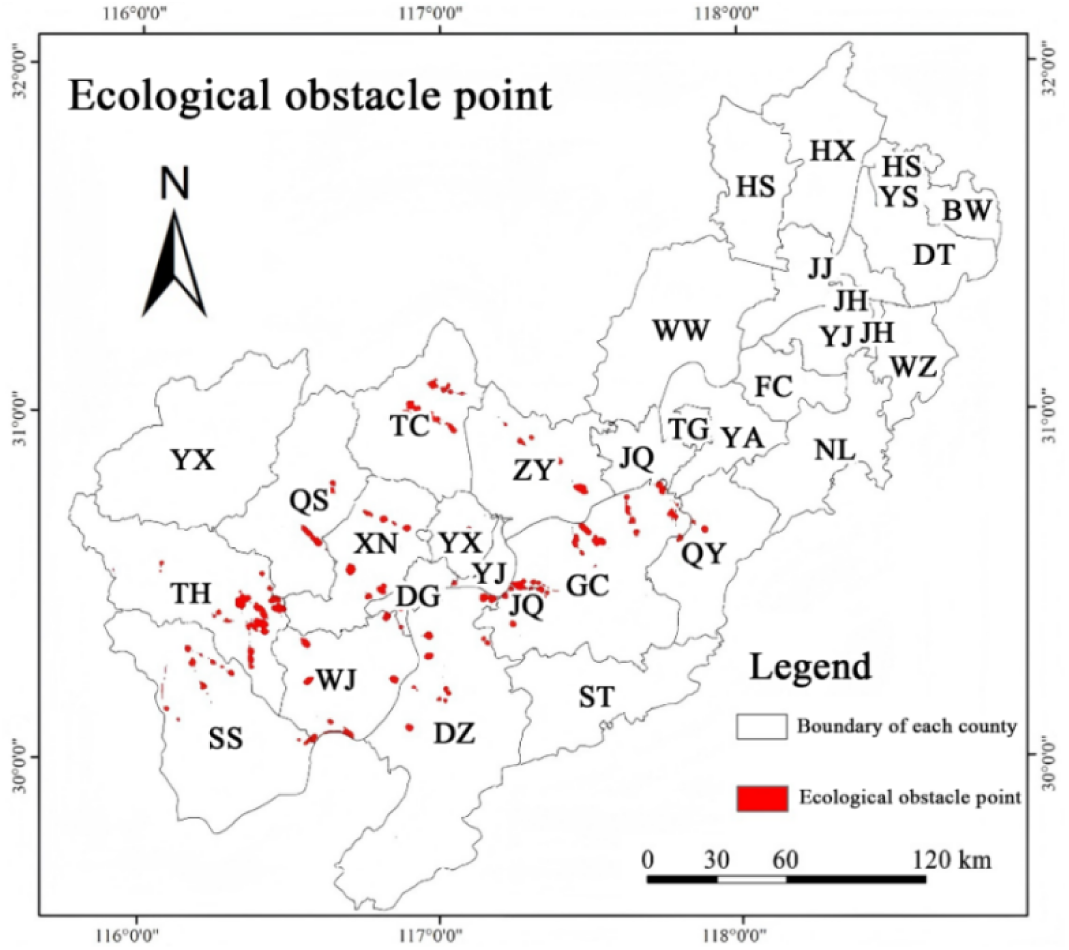
The distribution map of ecological obstacle points.

### 3.4 Discrimination of ecological warning points based on PLUS model

#### 3.4.1 Land use simulation in 2032

The 2032 land use simulation indicates a decline in cropland, grassland, and forest land, a slight increase in wetland and water area, and a pronounced expansion of construction land, which is projected to reach 4.21% of the total area. Construction land expansion is mainly concentrated around central urban areas and gradually extends toward surrounding regions. Spatially, expansion in the east–west direction is centered on Anqing, while north–south expansion is dominated by Ma’anshan and Wuhu. Urban expansion is particularly evident in Wuhu and Ma’anshan, where construction land encroaches upon cropland and grassland and progressively extends toward the plains of the Anhui section of the Yangtze River Basin. Owing to the predominantly flat terrain and the high proportion of cropland, land conversion to construction land is relatively favorable in this region. In contrast, the extensive ecological source areas in the western and southeastern parts of the study area effectively constrain construction land expansion. In the northern region, where ecological sources are scarce and land cover is dominated by wetland and water area, construction land expansion is more pronounced (Fig. 10).

**Fig. 10.**
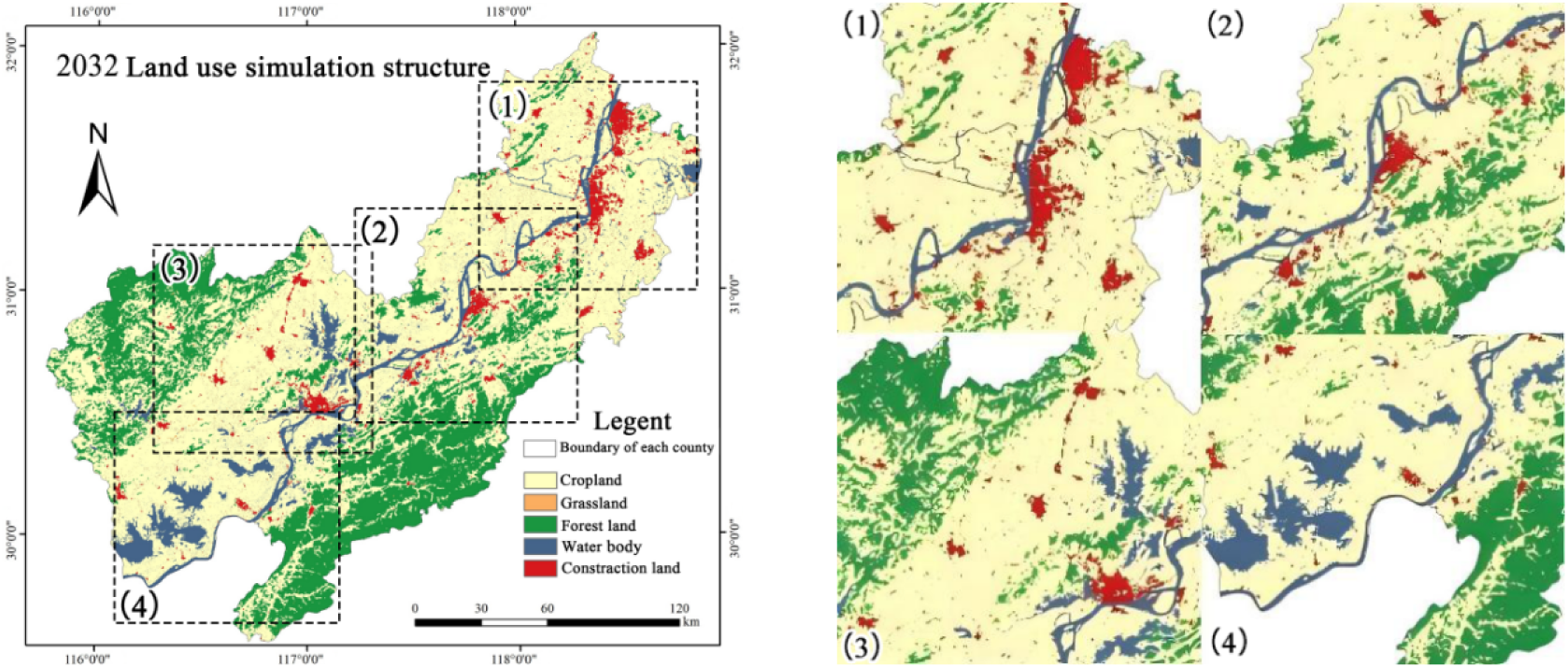
Land use distribution map and construction land expansion area in 2032.

#### 3.4.2 Distribution characteristics of ecological warning points

As shown in Fig. 11(a), severely sensitive areas within construction land expansion hotspots exhibit pronounced spatial heterogeneity, forming a clear “near-strong and far-weak” relationship with the Yangtze River shoreline. These areas are mainly concentrated in Ma’anshan, Wuhu, Tongling, and the urban districts of Anqing and Chizhou, where they significantly impede ecological connectivity between western and southeastern ecological sources. Highly sensitive areas show clear patterns of outward expansion and spatial aggregation around severely sensitive zones and tend to extend toward key ecological corridors linking the Dabie Mountains and Jiuhua Mountains, particularly in Susong of Anqing. In some locations, highly sensitive areas overlap directly with ecological sources in the southwestern part of the river basin, indicating elevated ecological risk. By overlaying construction land expansion hotspots with ecological sources and corridors, a total of 17 ecological warning points were identified in the study area (Fig. 11b), covering an area of 4,341.18 km². These warning points are dominated by cropland, which accounts for 65.06% of the total area, followed by construction land (20.96%) and wetland and water area (10.82%), while grassland represents only a small proportion (0.50%). The spatial distribution of ecological warning points indicates that ecological spaces in several cities, including Ma’anshan, Wuhu, and Tongling, are highly vulnerable to the expanding ecological stress effects of construction land. In particular, construction land expansion in urban districts blocks key ecological corridors, posing a substantial threat to the integrity of the regional ecological security pattern, such as Jinghu of Wuhu, Huashan of Ma ’anshan, Yushan, Yixiu District and Yingjiang of Anqing City.

**Fig. 11.**
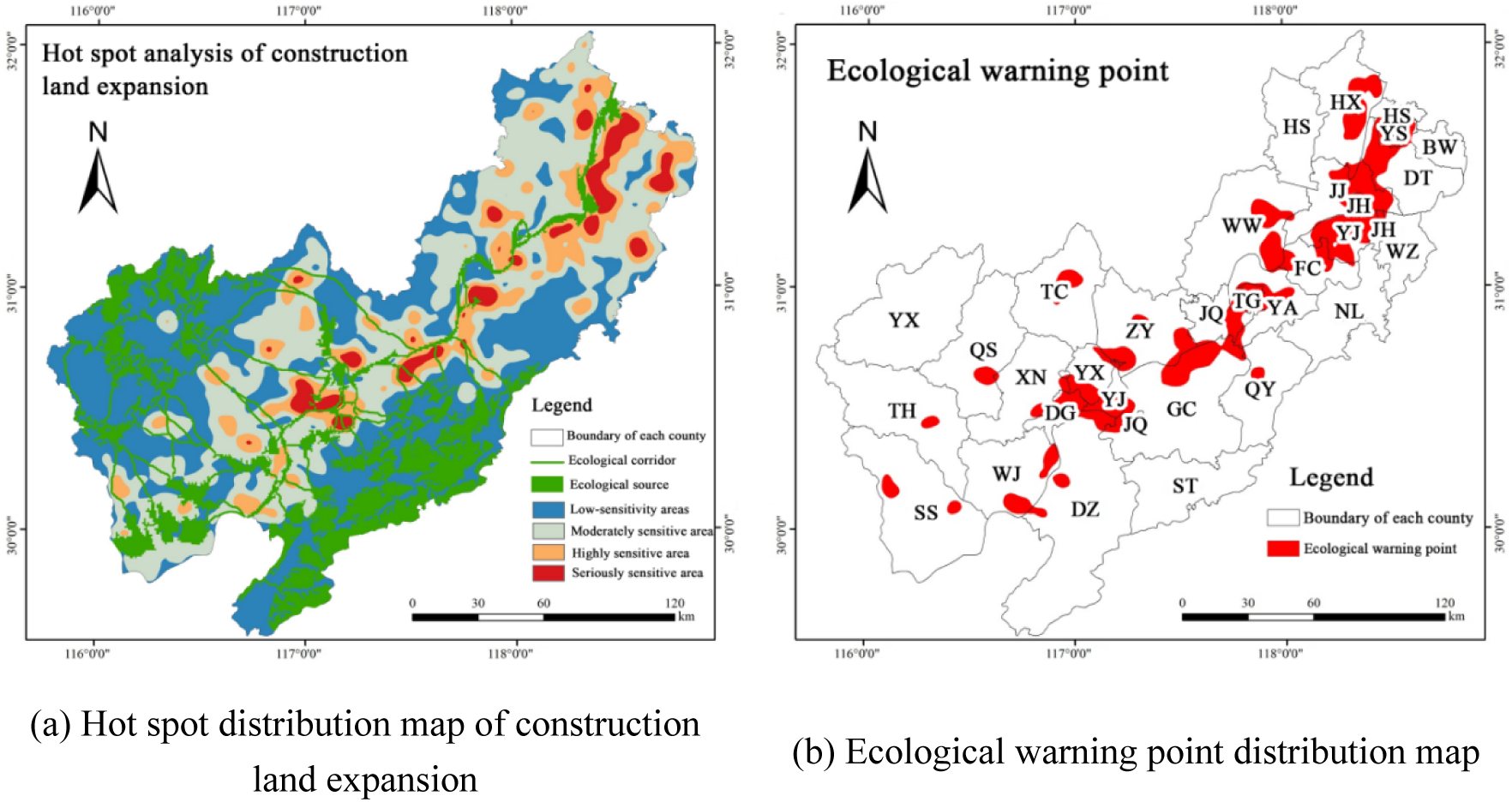
Hot spot analysis map of construction land expansion and distribution of ecological warning points in the study area from 2022 to 2032

#### 3.4.3 Ecological restoration zoning

Guided by natural geographical units and ecological processes, the study area was divided into four ecological restoration zones (Fig. 12). Areas where ecological sources, corridors, and pinch points overlap were designated as priority protection zones, covering 7,339.59 km² (21.02% of the study area) and dominated by forest land, wetland, and water area, serving as core ecological barriers. Ecological obstacle point areas were classified as core ecological restoration zones, covering 515.81 km² and mainly distributed in the southeastern Dabie Mountains and the Yangtze River alluvial plain, where cropland predominates. Severely and highly sensitive areas within construction land expansion from 2022 to 2032 were identified as moderate urban development zones, covering 5,401.44 km² and mainly located along the Yangtze River. Areas where ecological sources, ecological pinch points, ecological obstacle points, and ecological warning points intersect were defined as ecological space conflict zones, covering 213.58 km², characterized by high fragmentation and dominated by cropland and construction land.

**Fig. 12.**
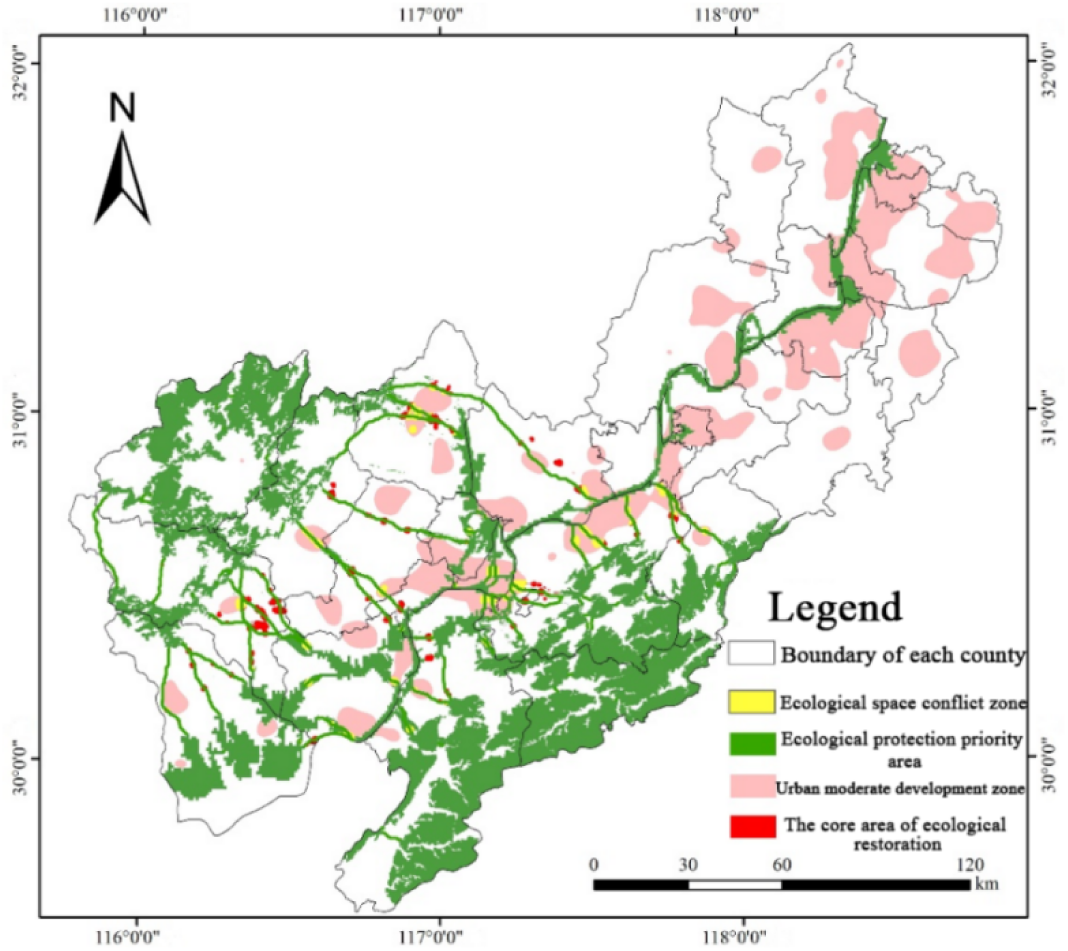
Ecological restoration zoning map of the study area.

## 4. Discussion

### 4.1 Implications for future urban and ecological planning of the Anhui section of the Yangtze River Basin

The results of this study extend existing ecological protection strategies by establishing a macro–meso–micro framework for ecological pattern planning in 2032. This systematic approach enables a multi-scale assessment of ecosystem structure and function, thereby providing a scientific basis for optimizing spatial configuration and implementing targeted ecological protection. Specifically, the framework is implemented through the following steps:

At the macro level, a multidimensional framework of “ecological background prioritization–functional trade-off regulation–ecological network optimization” was established. By integrating ecological vulnerability assessment and ecosystem service importance evaluation, differentiated ecological restoration zoning was delineated for the Anhui section of the Yangtze River Basin. In addition, ecological warning points predicted by the PLUS model were incorporated to construct the ecological planning pattern for 2032, characterized as “two belts, three zones, and multiple points” (Fig. 13). The “two belts” include the Yangtze River ecological protection belt and the Dabie–Jiuhua Mountains ecological corridor belt, which surround core areas and are dominated by forest land, wetland, and water area. These belts constitute a critical component of the regional ecological security pattern and function as buffers that enhance landscape connectivity. The “multiple points” refer to key ecological protection and restoration areas with high ecological importance, reflecting the region’s ecological background of rich biodiversity, dense water networks, and prominent ecological barrier functions.

**Fig. 13.**
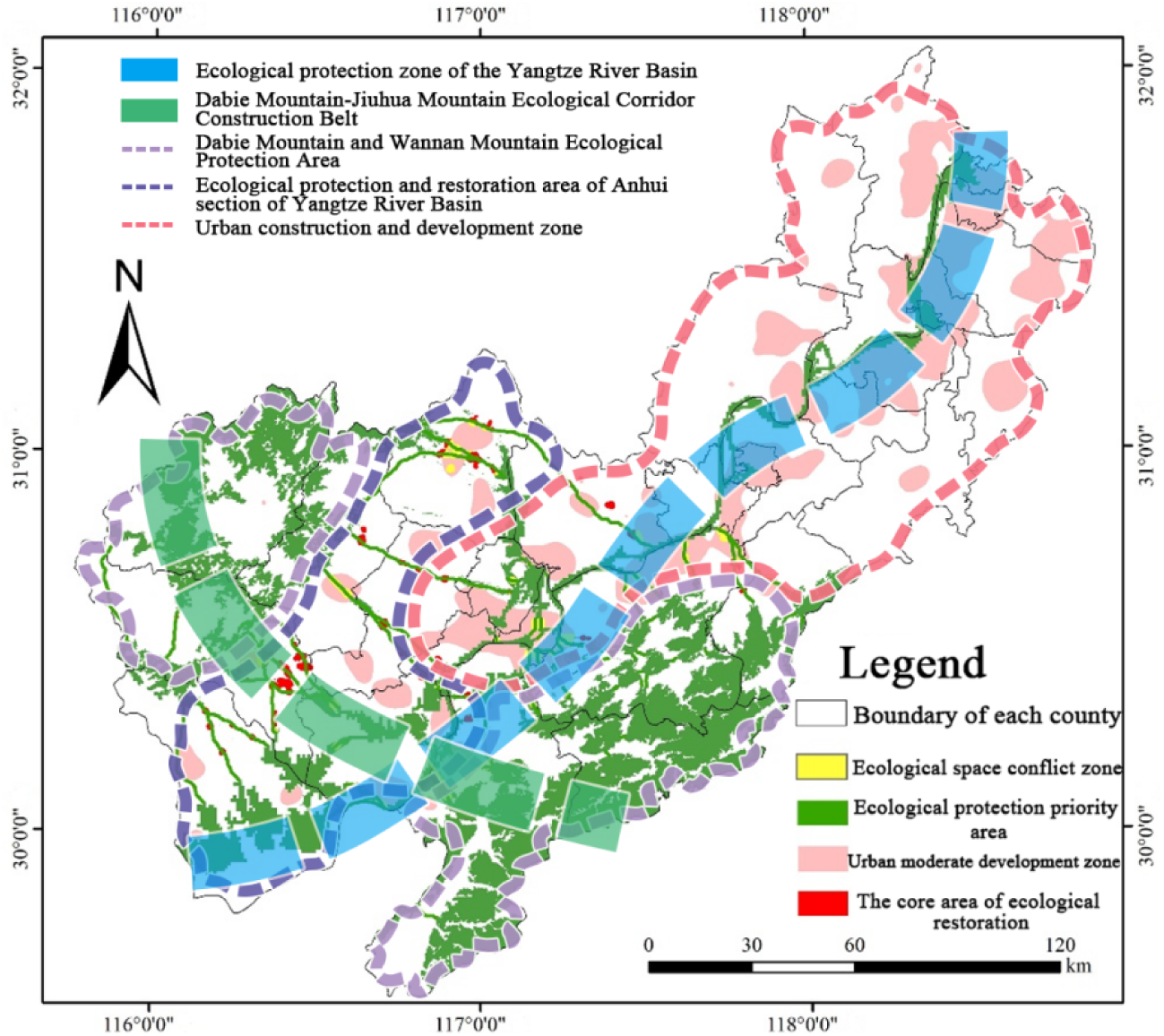
Ecological Planning Map of Anhui Section of the Yangtze River Basin in 2032.

At the meso level, regional ecological restoration zones for 2032 were precisely delineated based on ecological security zones, ecological pinch points, obstacle points, and ecological warning points, and corresponding regional control measures were proposed. By transcending administrative boundaries, this approach supports integrated planning and construction of ecological corridors, mitigates landscape fragmentation, and promotes coordinated development between urban systems and the natural environment. MSPA and landscape connectivity analyses were further applied to refine the classification and importance assessment of ecological regions. The resulting pattern of ecological protection importance is jointly shaped by the Yangtze River system, mountain forests, and wetland ecosystems, exhibiting a spatial configuration characterized by “river-oriented continuity, mountain dominance, wetland mosaics, and urban fragmentation”.

The results indicate that areas of extremely high ecological protection importance are mainly concentrated in the urban belt along the Yangtze River, dominated by wetland and water area, while highly important areas are primarily distributed in the southern part of the study area and are mainly composed of forest land. In this study, ecological protection importance increases with ecological vulnerability, which is consistent with the findings of Fang Chen et al. (plus literature). However, discrepancies among studies may arise from differences in the selection of ecosystem service function categories. This study further confirms that areas with higher ecosystem service function values tend to exhibit higher ecological protection importance (plus literature). The findings are also consistent with Wei Chenbo et al., who reported that extremely important and highly important areas are largely associated with forest land and water areas (plus literature). Variations in evaluation results across studies can be attributed to differences in service function indicators, weight assignment, and data sources. Future research should further explore the coupling relationships among ecosystem service functions, ecological vulnerability, and sensitivity, optimize weighting methods, and integrate multi-source data to enhance the scientific robustness and practical applicability of ecological protection importance assessments.

The results indicate that areas of extremely high ecological protection importance are mainly concentrated in the urban belt along the Yangtze River, dominated by wetland and water area, while highly important areas are primarily distributed in the southern part of the study area and are mainly composed of forest land. In this study, ecological protection importance increases with ecological vulnerability, which is consistent with the findings of Fang Chen et al. (45). However, discrepancies among studies may arise from differences in the selection of ecosystem service function categories. This study further confirms that areas with higher ecosystem service function values tend to exhibit higher ecological protection importance. The findings are also consistent with Wei Chenbo et al., who reported that extremely important and highly important areas are largely associated with forest land and water area (46). Variations in evaluation results across studies can be attributed to differences in service function indicators, weight assignment, and data sources. Future research should further explore the coupling relationships among ecosystem service functions, ecological vulnerability, and sensitivity, optimize weighting methods, and integrate multi-source data to enhance the scientific robustness and practical applicability of ecological protection importance assessments.

At the micro level, ecological planning and restoration for 2032 emphasize differentiated management to achieve ecological protection, low-carbon transition, and high-quality coordinated socio-economic development. Priority ecological protection areas are mainly located in the Dabie Mountain and the mountainou regions of southern Anhui, where forest land, wetland, and water area dominate. Wetland and water system in the Dabie Mountain, Jiuhua Mountain, and the Yangtze River Basin are incorporated into the ecological protection red line, and core biodiversity conservation zones are delineated. In these areas, human disturbance is strictly restricted, and measures such as hillside closure for natural regeneration, conversion of cropland to forest, slope restoration, and artificial wetland rehabilitation are implemented to enhance forest resistance, restore wetland function, and reduce anthropogenic impact. Core ecological restoration areas are mainly composed of ecological obstacle zones, dominated by forest land and cropland, where targeted restoration interventions are required.

Soil and water conservation should be strengthened in low mountain and hilly areas to control soil erosion, while soil improvement and crop rotation should be implemented in alluvial plains. Ecological buffer zones should be established to reduce agricultural non-point source pollution, and bioremediation and water purification technologies should be applied in mining areas to restore soil and water systems. Moderate urban development areas are mainly dominated by cropland and construction land, where strict cropland protection, improved land-use efficiency, and green urbanization should be promoted to balance economic growth and ecological protection. Measures including sustainable agricultural development, sponge city construction, water environment governance, urban–rural integration, green building promotion, and rural industry development should be advanced. At the institutional level, urban spatial structure optimization and the implementation of an “ecological compensation + green finance” mechanism are encouraged to promote coordinated ecological and economic development.

### 4.2 Advantages and rationality of research methods

In this study, the importance of ecosystem service functions and ecological vulnerability assessment results were integrated to identify key ecological protection areas in the Anhui section of the Yangtze River Basin. The InVEST model, SRP model, SPCA, and the maximum value method were jointly applied to evaluate ecological protection importance and delineate ecological sources. Ecological source identification comprehensively considered ecological protection importance, morphological spatial pattern analysis (MSPA), and landscape connectivity, enabling a multi-dimensional and robust assessment of potential ecological sources (47, 48). The extracted ecological sources are highly consistent with the major ecological barrier pattern proposed in *the Anhui Provincial Territorial Spatial Planning* (*2021–2035*) (49), and show a strong spatial overlap with the provincial ecological protection red line and *the 14th Five-Year Plan for Ecological Environment Protection of Anhui Province* (50), as well as the designated ecological priority protection areas in Anhui Province, demonstrating the scientific reliability and planning applicability of the proposed method (51).

The core ecological sources identified in this study cover most provincial nature reserves within the study area, including Shengjin Lake, Caizi Lake, and the Anqing Riverside Wetland Reserve. The comprehensive resistance surface was constructed by integrating natural conditions and human disturbance factors, and SPCA was applied to determine factor weights, ensuring the scientific robustness and overall representativeness of the resistance surface. This approach is consistent with the resistance surface construction method proposed by Deng Xin et al. (52). For ecological corridor identification, circuit theory was employed, which aligns with the methodology used by Zhang et al. (53). However, unlike previous studies that relied on empirical corridor width settings, this study further optimized corridor width by jointly analyzing cumulative current density responses and the proportion of non-ecological land under different thresholds. The results indicate that a corridor width of 30–100 m effectively maintains ecological connectivity while balancing ecological protection objectives and land-use constraints.

Previous studies have mainly focused on identifying existing ecological pinch points and obstacle points (54), but they are limited in revealing potential ecological risks under future land-use change. To address this limitation, this study introduces the PLUS model to dynamically simulate land-use patterns in 2032 and to quantify construction land expansion, thereby extending key area identification to include ecological warning points and forming an integrated identification framework. By comprehensively considering both natural and socioeconomic driving factors, the PLUS model improves the accuracy of regional risk identification (55) and supports the delineation of differentiated ecological planning zones for 2032. This approach facilitates the transition from static pattern analysis to dynamic risk-oriented planning, providing practical guidance for ecological restoration and the long-term optimization of the ecological security pattern in the Anhui section of the Yangtze River Basin.

### 4.3 Limitations and future research directions

This study constructs an ecological security pattern to identify differentiated ecological restoration zones and predict the ecological planning pattern of the Anhui section of the Yangtze River Basin in 2032. However, several limitations should be acknowledged. First, the ecological vulnerability and resistance assessment does not fully capture the heterogeneity of ecosystem types and biological communities. Future studies should incorporate long-term monitoring, high-resolution remote sensing, and field observations to improve evaluation accuracy. Second, although a suitable ecological corridor width is proposed for the study area, corridor width thresholds are highly region- and species-dependent, limiting their general applicability. Integrating multi-scale species movement data and ecological simulation models will be essential for refining corridor width determination in future research. Third, although the PLUS model combined with kernel density analysis effectively captures the potential ecological risks associated with construction land expansion, the large spatial extent of early warning areas limits fine-scale zoning and constrains differentiated management. Future studies should incorporate higher-resolution spatial analysis and multi-scenario simulations to improve the spatial precision of ecological warning point identification. Fourth, the identification of differentiated restoration areas based on circuit theory is highly sensitive to the resistance surface. Despite considering regional characteristics, some subjectivity remains in indicator selection and resistance assignment. Future research should account for the heterogeneous responses of different ecosystem types and biological communities by integrating long-term monitoring data, remote sensing inversion, and field surveys to enhance objectivity and accuracy. In addition, climate change effects are only partially considered in this study. Incorporating climate variability and future climate scenarios into ecological security pattern construction and restoration zoning will further improve the robustness and applicability of ecological planning outcomes.

## 5. Conclusions

Based on the integrated land-space elements of the Anhui section of the Yangtze River Basin, this study combines ecosystem service importance and ecological vulnerability assessments with multi-dimensional methods including MSPA pattern analysis, landscape connectivity analysis, and circuit theory. The suitable width of ecological corridors is comprehensively determined, a regional ecological security pattern is constructed, and the ecological planning pattern for 2032 is predicted. The main conclusions are as follows:

(1) The ecological protection importance pattern shows a “two mountains and one river” structure, with extremely and highly important areas accounting for about 25% of the study area. Extremely important areas are concentrated along the Yangtze River and dominated by wetland and water area, whereas highly important areas are mainly forested regions in the Dabie Mountain and Jiuhua Mountain.

(2) Thirty-seven ecological sources (6,901.39 km²) were identified, mainly in the Dabie Mountain and Jiuhua Mountain and the Yangtze River basin, dominated by forest land, wetland, and water area. Eighty-four ecological corridors (916.80 km) form a well-connected network, and a corridor width of 30–100 m is recommended.

(3) Ecological pinch points, obstacle points, and warning points were identified, covering 419.64 km², 515.81 km², and 4,341.18 km², respectively. Ecological pinch points mainly overlap with ecological corridors, while ecological obstacle points are primarily located at the edges of ecological source areas and are dominated by cropland. Based on the PLUS model, 17 ecological warning points were predicted for 2032, with a total area of 4,341.18 km², mainly consisting of cropland, followed by construction land and wetland and water area.

(4) The ecological planning pattern of the study area for 2032 was predicted, and four functional zones were delineated: ecological protection priority areas, core ecological restoration areas, urban moderate development areas, and ecological space conflict areas. Differentiated zoning management strategies were proposed at macro, meso, and micro levels, providing scientific guidance for identifying restoration priorities, formulating restoration plans, and selecting appropriate restoration measures. The results offer a transferable and reliable reference for regions facing similar ecological challenge.

## Author statement

Ruonan Li, Yuhan Xia and Shanshan Liang designed the experiments, Ruonan Li, Xia Yuhan and Shanshan Liang carried out the experiments, Yuhan Xia and Shanshan Liang wrote the manuscript. Wenjing Zhao, Mi Ping, Ziyu Chen and Yiwen Fan put forward relevant improvement suggestions.

## Acknowledgments

This work was supported by the National Natural Science Foundation of China [No:32371935 and 32371936], the Anhui Hefei Urban Ecosystem Research Station, National Forestry and Grassland Administration, Changjiang West Road 130, Shushan District, Hefei 230036, China, the project from the Key Laboratory of Tropical Island Land Surface processes and Environmental Changes of Hainan Province [DLZDSYS202404], and the Science and Technology Innovation Foundation of Key Lab of Biodiversity Conservation and Characteristic Resource Utilization in Southwest Anhui [Wxn202428].

## Data availability statement

Anyone who needs relevant data can contact the corresponding author.

